# Transcriptional variation and divergence of host-finding behaviour in *Steinernema carpocapsae* infective juveniles

**DOI:** 10.1101/272641

**Authors:** Neil D. Warnock, Deborah Cox, Ciaran McCoy, Robert Morris, Johnathan J. Dalzell

## Abstract

*Steinernema carpocapsae* is an entomopathogenic nematode that employs nictation and jumping behaviours to find host insects. We aimed to investigate the transcriptional basis of variant host-finding behaviours in the infective juvenile (IJ) stage of three *S. carpocapsae* strains (ALL, Breton and UK1). RNA-seq analysis revealed that whilst up to 28% of the *S. carpocapsae* transcriptome was differentially expressed (P<0.0001) between strains, remarkably few of the most highly differentially expressed genes (>2 log2 fold change, P<0.0001) were from neuronal gene families. *S. carpocapsae* Breton displays increased chemotaxis toward the laboratory host *Galleria mellonella*, relative to the other strains. This correlates with the up-regulation of four srsx chemosensory GPCR genes, and a sodium transporter gene, *asic-2*, relative to both ALL and UK1 strains. The UK1 strain exhibits a decreased nictation phenotype relative to ALL and Breton strains, which correlates with co-ordinate up-regulation of neuropeptide like protein 36 (*nlp-36*), and down-regulation of an srt family GPCR gene, and a distinct *asic-2-like* sodium channel paralogue. To further investigate the link between transcriptional regulation and behavioural variation, we sequenced microRNAs across IJs of each strain. We have identified 283 high confidence microRNA genes, yielding 321 isomiR variants in *S. carpocapsae*, and find that up to 36% of microRNAs are differentially expressed (P<0.0001) between strains. Many of the most highly differentially expressed microRNAs (>2 log2 fold, P<0.0001) are predicted to regulate a variety of neuronal genes that may contribute to variant host-finding behaviours. We have also found evidence for differential gene isoform usage between strains, which alters predicted microRNA interactions, and could contribute to the diversification of behaviour. These data provide deeper insight to the transcriptional landscape of behavioural variation in *S. carpocapsae*, underpinning efforts to functionally dissect the parasite host-finding apparatus.

**Author summary:** *Steinernema carpocapsae* is a lethal parasite of insects. In order to find and invade a host insect, the *S. carpocapsae* infective juvenile will typically stand upright, waving its anterior in the air as it searches for host-specific cues. When the infective juvenile senses insect volatile compounds and movement (both signals are required), it will attempt to jump towards the source of those stimuli. Whilst the jumping behaviour is unique to *Steinernema* species nematodes, nictation is a host-finding behaviour shared with other important parasites of medical and veterinary importance. We have found that different strains of *S. carpocpsae* use modified host-finding strategies, and that these behavioural differences correlate with gene expression patterns, identifying genes that may be crucial in regulating aspects of host-finding. We also assessed the complement of microRNAs, which are small non-coding RNAs that regulate target gene expression. We found a surprising difference in the abundance of shared microRNAs between strains of *S. carpocapsae;* these differences also reveal expression differences that correlate with behavioural variation. Predicted microRNA target genes suggest that microRNA variation could significantly influence the behaviour of nematodes. Broadly, this study provides insight to the relationship between gene expression and behaviour, paving the way for detailed studies on gene function.

## Introduction

Parasitic nematodes employ a variety of behaviours that maximise opportunity for host contact and invasion. These behaviours vary across species, ranging from passive reliance on host ingestion, through pro-active host-finding by migration, nictation, and even jumping [1–4]. Each of these behavioural strategies rely on the incorporation of multiple sensory inputs, spanning chemosensory, olfactory, mechanosensory, thermosensory and hygrosensory circuits [2, 5]. Nematode host-finding strategies are also remarkably plastic, varying in response to experience and environment [6]. Despite the obvious importance of parasite host-finding behaviour to medical, veterinary and agricultural interests, we know relatively little about the genes involved in regulating these behaviours. It may be possible to develop new approaches to parasite control by targeting components of the parasite host-finding apparatus.

The evident complexity of nematode behaviour belies their relative neuroanatomical simplicity. It is thought that the neurochemical complexity of nematodes is central to their diverse behavioural capacity and adaptability [7]; neuropeptides in particular are highly enriched and conserved between nematode species with diverse life history traits [8]. It has also been demonstrated that FMRFamide-like peptide (*flp*) genes are co-ordinately up-regulated in host-finding stages of diverse parasitic nematodes [9], and that they contribute to various host-finding behaviours [9, 10], along with insulin-like peptides [11] and neuropeptide-like proteins [12]. Diverse neuronal gene families also contribute to the surprising behavioural enrichment of such simple organisms [13]. Variation in gene transcript abundance and sequence identity is central to the phenotypic plasticity of cells, tissues and organisms, underpinning behavioural variation. Differential expression or isoform variation can be regulated by transcription factors in response to developmentally encoded programs, as well as in response to environmental input. Small non-coding RNAs also contribute to gene regulation, as a factor of developmental stage, and in response to environmental challenge. Small RNAs have been implicated in driving phenotypic novelty and adaptation within and between species [14 – 17].

MicroRNAs are small non-coding RNAs that negatively regulate target gene expression across higher organisms [18], and have been shown to modulate neuronal connectivity, synaptic remodelling [19] and memory within the olfactory system of other invertebrates [20]. *In vivo* cross-linking and immunoprecipitation of microRNA-specific argonautes coupled with sequencing of bound mRNA transcripts in *Caenorhabditis elegans*, demonstrates an enrichment of neuronal gene families. This confirms that many neuronal genes with known involvement in behaviour are biologically relevant microRNA targets [21]. Key to understanding the contribution of neuronal gene function and microRNA regulation to host-finding behaviour in parasitic nematodes, is the development of a suitable model system through provision of foundational datasets.

*Steinernema* spp. nematodes are obligate entomopathogens that invade and kill insect hosts through coordinated action with commensal *Xenorhabdus* bacteria [22]. *Steinernema* infective juveniles (IJs) display qualitatively different host-finding strategies between species [23], representing a unique resource for the comparative analysis of behaviour. *Steinernema carpocapsae* is generally considered to employ an ‘ambushing’ strategy, characterised by nictation and jumping behaviours. Nictation is enacted by nematodes that stand upright, waving their anterior in the air [24]. During nictation, the nematode can respond to sensory stimuli in one of three ways: (i) it can cease nictation and transition to a migratory phase; (ii) it can engage in a torpid ‘standing’ phenotype that may enhance opportunity for host attachment; and (iii) it may jump directionally in response to volatile and mechanosensory input [9, 24]. Whilst the jumping behaviour is thought to be unique to a small number of *Steinernema* spp., nictation is shared amongst many economically important animal parasitic nematodes, alongside the model nematode *C. elegans*, for which nictation represents a long-range phoretic dispersal behaviour [25]. In this study our aim was to profile the host-finding behaviours of *S. carpocapsae* strains, and to begin mapping the transcriptional landscape of behavioural variation across protein-coding and non-coding RNAs. The identification of genes or gene isoforms that are differentially expressed and which correlate with distinct behavioural states could improve our understanding of genotype-phenotype linkages. This foundational knowledge will underpin efforts to further dissect the parasite host-finding apparatus through detailed functional studies.

## Materials and Methods

### *S. carpocapsae* culture

*Steinernema carpocapsae* strains (ALL, Breton and UK1) were maintained in *Galleria mellonella* at 23°C. IJs were collected by White trap [26] in a solution of Phosphate Buffered Saline (PBS). Freshly emerged IJs were used for each experiment. Individual biological replicates and RNAseq libraries described below were derived from a mixed population of IJs that emerged from multiple *G. mellonella* cadavers on the same day.

### Behavioural assays

*Galleria mellonella* host-finding assays and dispersal assays were conducted as published [9]; in both instances five biological replicates were assayed across three technical replicates each. For the nictation assays, micro-dirt chips were made from a PDMS mould [27], with 3.5% ddH2O agar. 20 IJs suspended in 1.5 µl PBS were pipetted onto the micro-dirt chip, under a binocular light microscope. Once the liquid had evaporated and the IJs could move freely, the number of nictating IJs was counted at 1, 2.5 and 5 minute intervals. Nictation assays were conducted over five biological replicates, each with five technical replicates of 20 IJs each. Each dataset was assessed by Brown-Forsythe and Bartlett’s tests to examine homogeneity of variance between groups. One way ANOVA and Tukey’s multiple comparison tests were then used to assess statistically significant differences in mean across experimental groups. Tests were conducted in Graphpad Prism 7.02.

### RNA-seq, differential expression and isoform variant analysis

Six biological replicates of each strain were prepared from approximately 10,000 individuals (80 µl packed volume after centrifugation at 2000 rpm for 2 minutes) each. Total RNA was extracted from IJs using TRIzol Reagent (Invitrogen) and DNase treated using the Turbo DNase kit (Ambion) following manufacturer’s instructions. RNA quantity were assessed by gel electrophoresis and quantified using Qubit RNA BR Assay Kit (Life Technologies) as per manufacturer’s instructions. A total of six transcriptome libraries (150 bp, paired end) were prepared for each *S. carpocapsae* strain, from 1 µg of total RNA each, using the TruSeq RNA Library Prep Kit v2 (Illumina) following manufacturer’s instructions. Sequencing was performed on the HiSeq2500 instrument. Fastq files were assessed for quality using the FastQC (v. 0.11.3) package [28]. Adapters, low quality bases, and reads shorter than 36 bp were removed using the Trimmomatic (v. 0.35) package [29]. The trimmed data quality was then re-assessed using the FastQC package. The *S. carpocapsae* genomic contigs (PRJNA202318.WBPS9) and associated GFF file (PRJNA202318.WBPS9) were downloaded from https://parasite.wormbase.org/index.html. Annotations were converted to GTF format using Cufflinks (v. 2.2.2.20150701) [30]. High quality reads were then mapped to the S. *carpocapsae* genome [31] using the STAR (v. 2.5.3a) package [32]. Isoform expression levels were quantified using the RSEM (v. 1.2.19) package [33], and integrated EBseq package [34]. Raw read counts mapping to each gene in each sample were consolidated into a single count table. This process was repeated for each isoform. Downstream analyses was performed using R (v. 3.3.1) [35]. Differential expression of genes was quantified using the DESeq2 (v. 1.14.1) package [36]. Graphics were generated in R using RColorBrewer (v. 1.1-2) [37], gplots (v. 3.0.1) [38], geneplotter (v. 1.52) [39] and pheatmap (v. 1.0.8) [40] packages with custom R scripts.

### MicroRNA sequencing, discovery and quantification

Six biological replicates of each strain were prepared from approximately 10,000 individuals (80 µL packed volume after centrifugation at 2000 rpm for 2 minutes) each. Total RNA was extracted from IJs using TRIzol Reagent (Invitrogen) and DNase treated using the Turbo DNase kit (Ambion) following manufacturer’s instructions. Small RNA libraries were generated from the same RNA samples that were used for matched transcriptomes. RNA quantity was assessed by gel electrophoresis and quantified using Qubit RNA BR Assay Kit (Life Technologies) as per manufacturer’s instructions. A total of six small RNA libraries were prepared for each strain, from 1 µg of total RNA each, using the TruSeq Small RNA library Kit (Illumina) following manufacturer’s instructions. 50 bp single-end libraries were sequenced on the HiSeq2500 instrument. Fastq files were assessed for quality using the FastQC (v. 0.11.3) package [28]. Adapters, low quality bases, and reads shorter than 13 bp were removed using the Cutadapt package (v. 1.8) [41]. Sequence reads without adapters were also discarded. Reads that passed QC were mapped to the genome sequence of *S. carpocapsae*, and microRNAs were identified by miRDeep2 (v. 2.0.0.8) [42], using a training set of mature and precursor microRNA sequences downloaded from miRBase (http://www.mirbase.org/). Naming of microRNAs was preferentially aligned with *C. elegans*, as indicated by miRDeep2 output. Novel *S. carpocapsae* microRNAs were named and numbered sequentially, taking care to avoid overlap with any *C. elegans* microRNA. Differentially expressed microRNAs were identified as above, using the DESeq2 package, and were presented using RColorBrewer, gplots, geneplotter and pheatmap packages.

### MicroRNA target gene prediction

Three and five prime UnTranslated Regions (UTRs) of computationally predicted *S. carpocapsae* genes were exported from wormbase parasite [31, 43] using the biomart function. Retrieved sequences were converted to fasta format and predicted microRNA binding sites were identified using miRanda [44]. Two separate miRanda analyses were performed, using i) unrestricted settings and ii) strict settings that require perfect conservation of seed site sequence complementarity between microRNA and target mRNA. Experimental verification of microRNA target predictions indicate that perfect complementarity between the microRNA seed region and target mRNA provides the highest degree of specificity and sensitivity [45]. However, Argonaute CLIP-seq analyses indicate that around 40% of all microRNA-mRNA interactions lack perfect seed region complementarity [46]; limiting analyses to perfect seed region requirements will lead to a substantial number of false negatives. MiRanda allows analyses that span canonical seed region complementarity, and non-canonical interactions, providing a robust overview of interactions that follow experimentally validated examples [45]. In each instance, we have included information of relative target site predictions using both strict and unrestricted target identification approaches.

### Annotation of neuropeptide, neurotransmitter, GPCR, innexin and ion channel genes

*S. carpocapsae* neuropeptide gene orthologues were identified via reciprocal BLAST analysis. A list of available *C. elegans* FLP, NLP and INS pre-propeptide sequences were obtained from Wormbase [47] and used as BLASTp and tBLASTn search strings via the Wormbase ParaSite BLAST server [43] under default settings. The protein sequences for overlapping genes associated with each high scoring pair were then employed as BLASTp search strings against the available *C. elegans* protein dataset via the Wormbase BLAST server [47]. Where no overlapping gene annotation was available, novel predicted proteins were generated by concatenating high-scoring return sequences to facilitate reciprocation [48]. The top reciprocal BLAST hit from *C. elegans* was used to assign putative neuropeptide gene names, and subsequent manual comparison of *S. carpocapsae* neuropeptide primary sequences to established nematode neuropeptide motifs [49] was used to confirm or reassign gene names where appropriate. GPCR, ion channel and innexin genes were exported from Wormbase parasite according to GO term, using the biomart function. All neurotransmitter genes were identified by reciprocal BLAST, as above. Additionally, the identity of each gene represented in heatmaps and tables of this manuscript was confirmed by reciprocal BLAST, as above. In a number of instances, different *S. carpocapsae* genes reciprocated to the same *C. elegans* gene. In any such case, a simple numbering system was applied to gene names in order to reflect clustered identity.

## Results

### Behavioural variation across *S. carpocapsae* strains

*S. carpocapsae* Breton demonstrated an increased chemotaxis index in response to *G. mellonella* volatiles, indicating that significantly more IJs migrated toward the larvae relative to other strains (Fig 1A; 0.88 ±0.009, relative to 0.43 ±0.06 for ALL, and 0.31 ±0.08 for UK1, P<0.0001****). *S. carpocapsae* UK1 exhibited a reduced nictation phenotype relative to the other strains (Fig 1B; 2.26% ±0.99 of UK1 IJs were nictating at the 5 minute time point, relative to 26.5% ±3.8 of ALL strain IJs, and 27.7% ±3.7 of Breton strain IJs, P<0.0001****). No statistically significant difference was observed in the dispersal behaviour of IJs from *S. carpocapsae* strains (Fig 1C).

**Figure 1.**
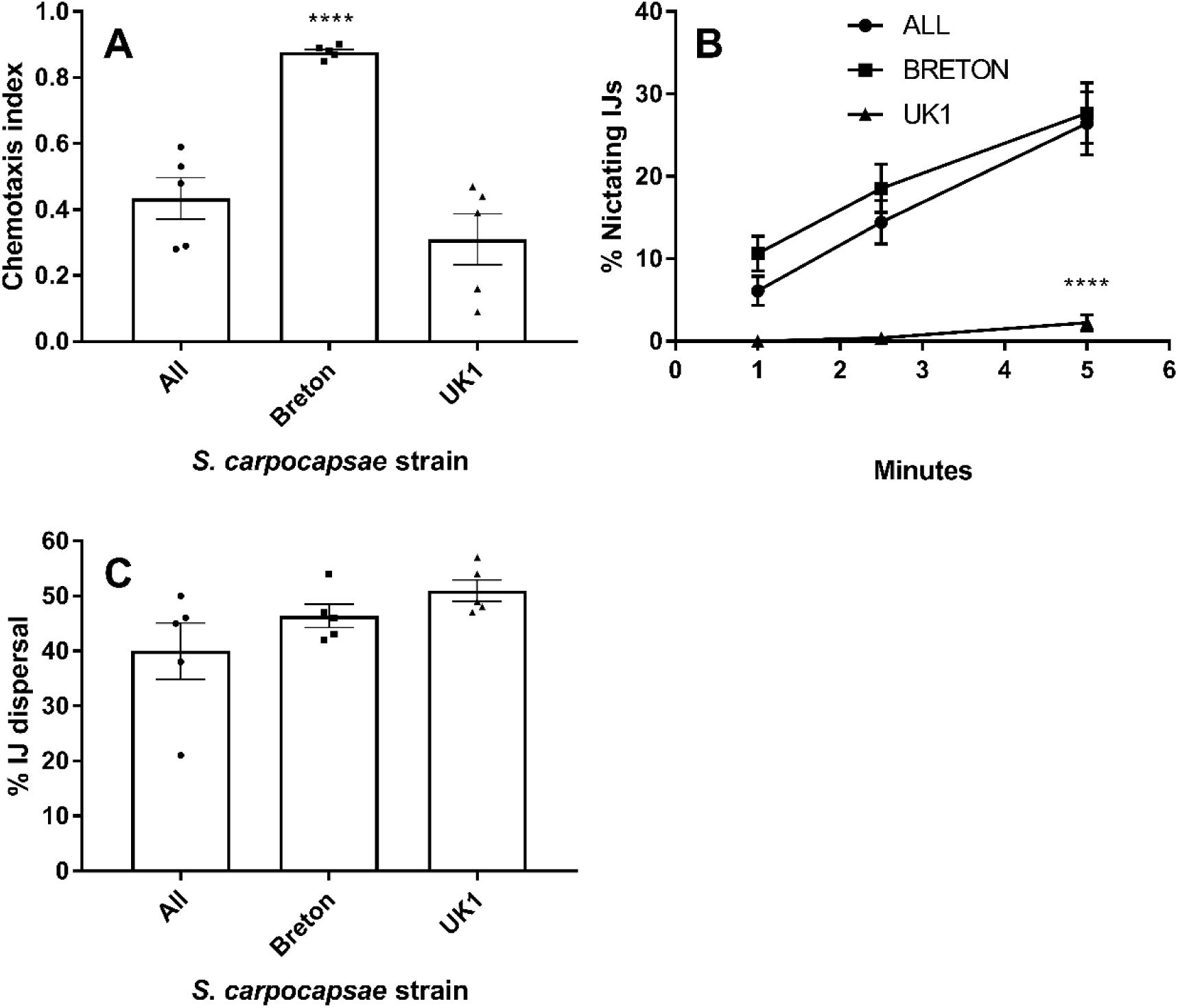
*S. carpocapsae* strains display variant host-finding behaviours. (A) Mean chemotaxis index of *S. carpocapsae* strains in response to *Galleria mellonella* larvae; (B) Mean number of nictating *S. carpocapsae* IJs over a time-course; (C) Mean percentage dispersal of *S. carpocapsae* IJs into the peripheral assay zone (dispersed) after a 1 hour timecourse. Data points represent mean ±SEM; assessed by ANOVA and Tukey’s multiple comparison test using Graphpad Prism 7.02; P<0.0001****.

### Transcriptional variation across strains

7,494 (28%) and 3,662 (13.7%) of *S. carpocapsae* UK1 genes were differentially expressed (P<0.0001****) relative to Breton and ALL strains, respectively (Fig 2; supplemental file Sz). 4,762 (17.7%) of *S. carpocapsae* Breton genes were differentially expressed (P<0.0001****) relative to the ALL strain (Fig 2). 22 (0.08%) of the 1,509 (5.3%) most highly differentially expressed genes across all pairwise strain comparisons (>2 log2 fold, P<0.0001****) were representative of neuronal gene families, based on GO term annotations (supplemental file S1). Of these 22 nueronal genes, only eight were observed to share differential expression patterns across pairwise comparisons that correlate with distinct IJ behaviours; either increased chemotaxis (*S. carpocapsae* Breton; Fig 1A) or decreased nictation (*S. carpocapsae* UK1; Fig 1B). Specifically, shared up or down-regulation in *S. carpocapsae* Breton relative to both ALL and UK1 strains would correlate gene differential expression with increased chemotaxis to *G. mellonella* (Fig 1A). Conversely, shared up or down-regulation in *S. carpocapsae* UK1 relative to both Breton and ALL strains would correlate gene differential expression with reduced nictation behaviour (Fig 1B). In order to explore the transcriptional regulation of neuronal gene families implicated in these behavioural differences, we assessed each gene family represented in the highly differentially expressed category, as defined above. This encompassed G-protein coupled receptors (GPCRs), ion channels, innexins, neuropeptides and neurotransmitter synthesis, degradation and transport genes (supplemental file S1).

**Figure 2.**
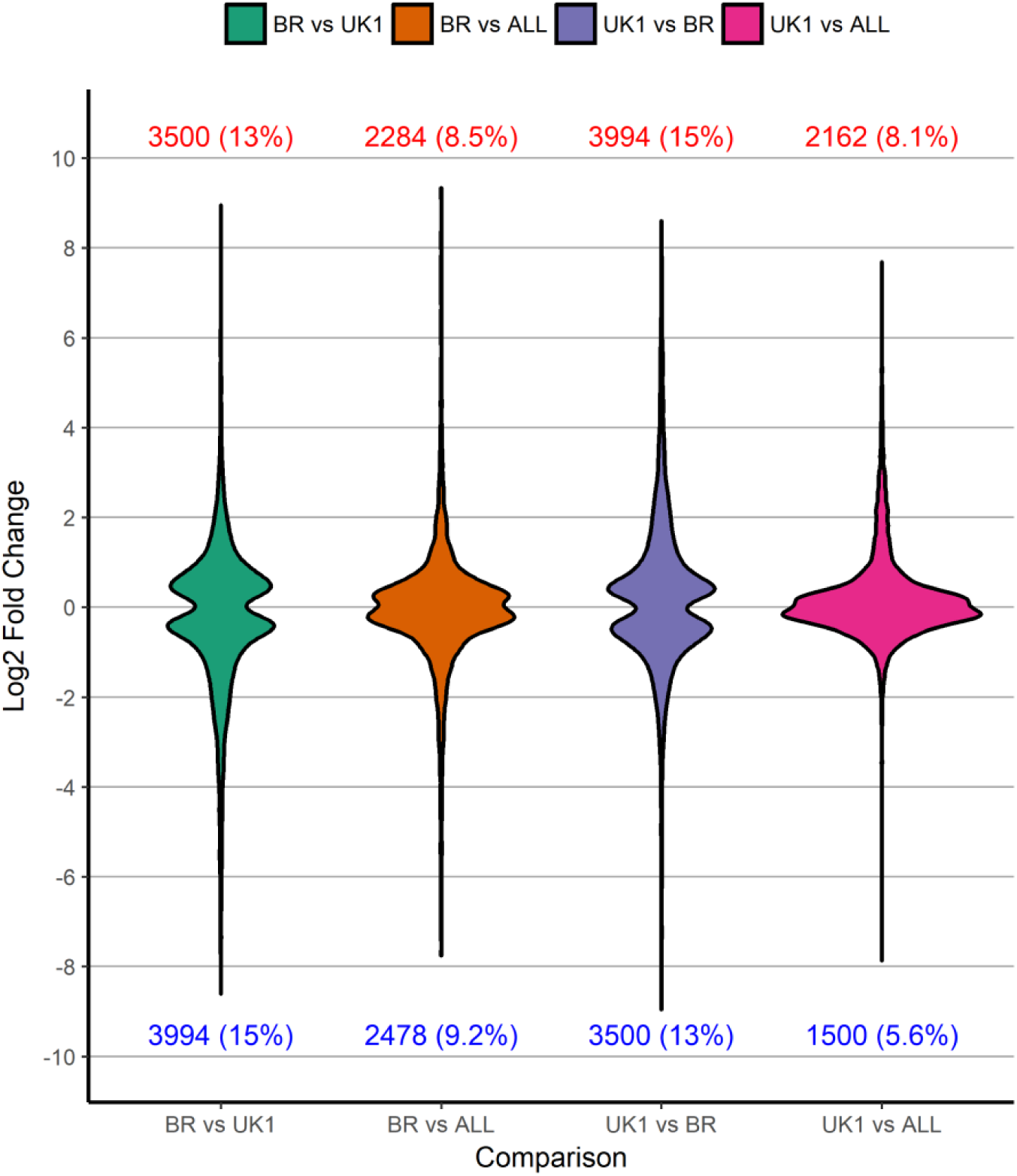
Violin plot showing significantly up-regulated and down-regulated genes across pairwise *S. carpocapsae* strain comparisons. Statistically significant (P<0.0001****) differentially expressed genes are plotted irrespective of log2 fold change. Total number and relative percentage of up-regulated and down-regulated genes are presented above and below each plot, respectively.

### Neuropeptide and neurotransmitter genes

*Nlp-36* was the only neuropeptide-like protein gene to demonstrate significant fold change differences (>2 log2 fold, P<0.0001****) that correlate with decreased nictation behaviour in *S. carpocapsae* UK1 across pairwise comparisons (up-regulated 2.8 and 3.5 log2 fold, P<0.0001**** relative to Breton and ALL strains, respectively) (Fig 3A-B; supplemental file S2). Though falling short of our shared >2 log2 fold change threshold, insulin-like peptide gene, *daf-28*, was the single most differentially regulated *ins* gene, exhibiting a polarised expression pattern that correlates inversely with both enhanced chemotaxis toward *G. mellonella* (*S. carpocapsae* Breton), and reduced nictation behaviour (*S. carpocapsae* UK1) (down-regulated 1.1 and 2.8 log2 fold, P<0.0001**** in Breton relative to ALL and UK1, respectively; up-regulated 1.7 and 2.8 log2 fold, P<0.0001**** in UK1 relative to ALL and Breton, respectively) (Fig 2. C-D; supplemental file S2). *Flp* genes were comparatively less variant between strains, with *flp-34* exhibiting the largest pairwise expression change that correlates with increased chemotaxis behaviour in *S. carpocapsae* Breton (up-regulated 0.7 and 0.6 log2 fold, P<0.0001****, relative to UK1 and ALL strains, respectively) (supplemental file S2). The tyrosine decarboxylase gene, *Sc-tdc-1*, was the most differentially expressed neurotransmitter gene (down-regulated 1.4 and 1.9 log2 fold, P<0.0001**** in *S. carpocapsae* UK1 relative to ALL and Breton, respectively), correlating with decreased nictation behaviour, but falling short of a shared >2 log2 fold change threshold (supplemental file S2).

**Figure 3.**
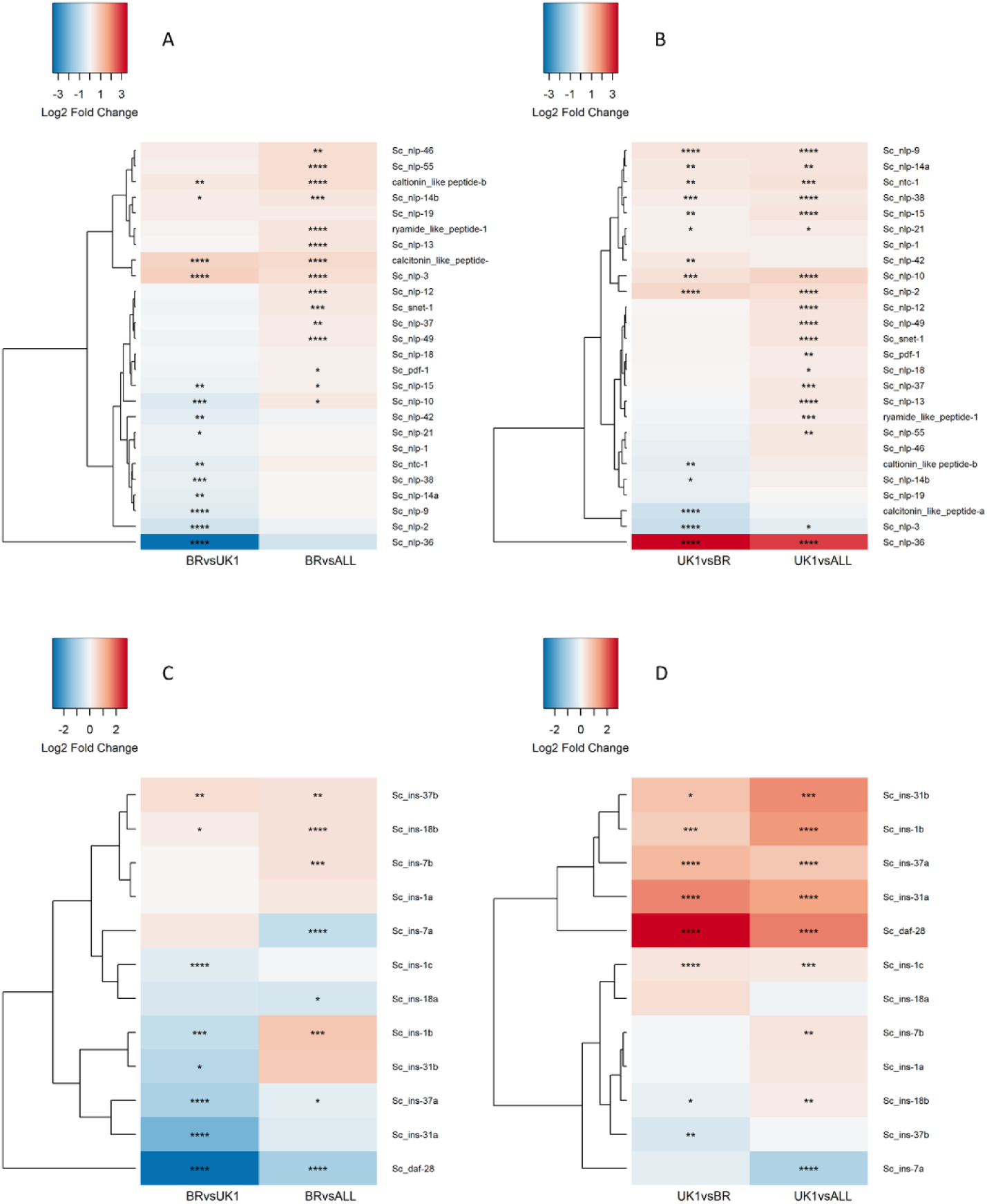
Differential expression analysis of neuropeptide gene families that correlate with *S. carpocapsae* strain behaviour. Figures A and C represent pairwise comparisons of *S. carpocapsae* Breton (BR) relative to UK1 and ALL; assessing shared gene expression patterns that correlate with increased attraction to *G. mellonella*; Figures B and D represent pairwise comparisons of *S. carpocapsae* UK1 relative to BR and ALL; assessing gene expression patterns that correlate with reduced nictation behaviour. (A-B) Differential expression analysis of neuropeptide-like protein genes; (C-D) differential expression of insulin-like peptide genes; adjusted P values are indicated for all log2 fold changes; padj P<0.05*. P<0.01**, P<0.001***, P<0.0001****. *Flp* and neurotransmitter gene maps are included in supplemental as there are no differentially expressed genes satisfying a >2 log2 fold threshold.

### GPCR, innexin and ion channel genes

Significant up-regulation of four srsx GPCR genes (*Sc-srsx-25v, Sc-srsx-3ii, Sc-srsx-22i*, and *Sc-srsx-24ii*) (>2 log2 fold, P<0.0001****) correlates with increased chemotaxis of *S. carpocapsae* Breton to the insect host *G. mellonella*, relative to UK1 and ALL strains (Fig 4A; supplemental file S2). By way of contrast, the *Sc-srt-62* chemosensory GPCR gene was co-ordinately down-regulated in *S. carpocapsae* UK1, correlating with reduced nictation behaviour (down-regulated 2.6 and 2.5 log2 fold, P<0.0001**** relative to ALL and Breton strains, respectively) (Fig 4B; supplemental file S2). Although falling below our threshold of shared >2 log2 fold change across both strain pairwise comparisons, *Sc-inx-7ii* demonstrated the largest expression change in the innexin / gap junction gene family, correlating with increased chemotaxis behaviour (down-regulated 1.03 and 2.65 log2 fold in *S. carpocapsae* Breton relative to UK1 and ALL strains, respectively; Fig 4C-D; supplemental file S2). Two paralogous *asic-2* sodium channel genes were found to be differentially expressed, and inversely related to altered chemotaxis behaviour (*Sc-asic-2ii*; down-regulated 2.3 and 2.7 log2 fold, P<0.0001**** in *S. carpocapsae* Breton, relative to ALL and UK1 strains, respectively; Fig 4E, supplemental file S2), and reduced nictation behaviour (*Sc-asic-2i*; up-regulated 2.9 and 3 log2 fold, P<0.0001**** in *S. carpocapsae* UK1, relative to ALL and Breton strains, respectively; Fig 4F, supplemental file S2).

**Figure 4.**
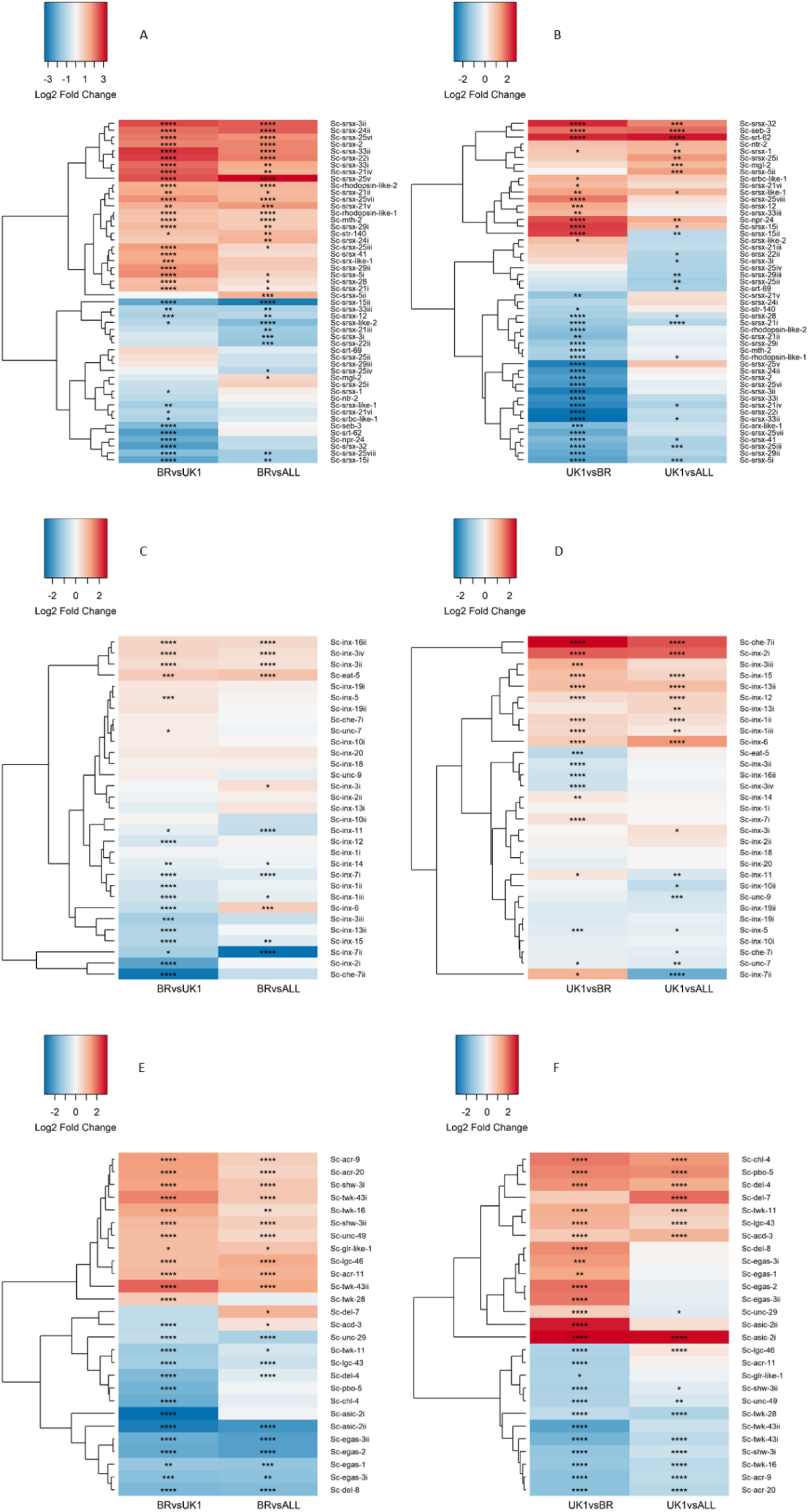
Differential expression analysis of GPCR, innexin and ion channel gene families that correlate with *S. carpocapsae* strain behaviour. Figures A, C and E represent pairwise comparisons of *S. carpocapsae* Breton (BR) relative to UK1 and All; assessing shared gene expression patterns that correlate with increased chemotaxis to *G. mellonella*; Figures B, D and F represent pairwise comparisons of *S. carpocapsae* UK1 relative to BR and All; assessing gene expression patterns that correlate with reduced nictation behaviour. (A - B) putative GPCR genes with >1 log2 fold difference between at least one of the pairwise comparisons; (C and D) putative innexin genes across all log2 fold changes; (E and F) putative ion channel genes with >1 log2 fold difference between at least one of the pairwise comparisons. P values are indicated for all log2 fold changes; padj P<0.05*. P<0.01**, P<0.001***, P<0.0001****.

### MicroRNA prediction and quantification

Following miRDeep2 identification, quality control and manual curation, a total of 283 high confidence (reads identified in at least five of the six libraries per strain) microRNA genes and 321 predicted isoMir variants were identified across a deep analysis (n=18 libraries) of *S. carpocapsae* strains (Supplemental file S3). 102 (36%) and 103 (36%) *S. carpocapsae* Breton microRNAs were differentially expressed (P<0.0001****) relative to UK1 and ALL strains, respectively. 50 (17.6%) *S. carpocapsae* UK1 microRNAs were differentially expressed (P<0.0001****) relative to *S. carpocapsae* ALL (Fig 5; supplemental file S3).

**Figure 5.**
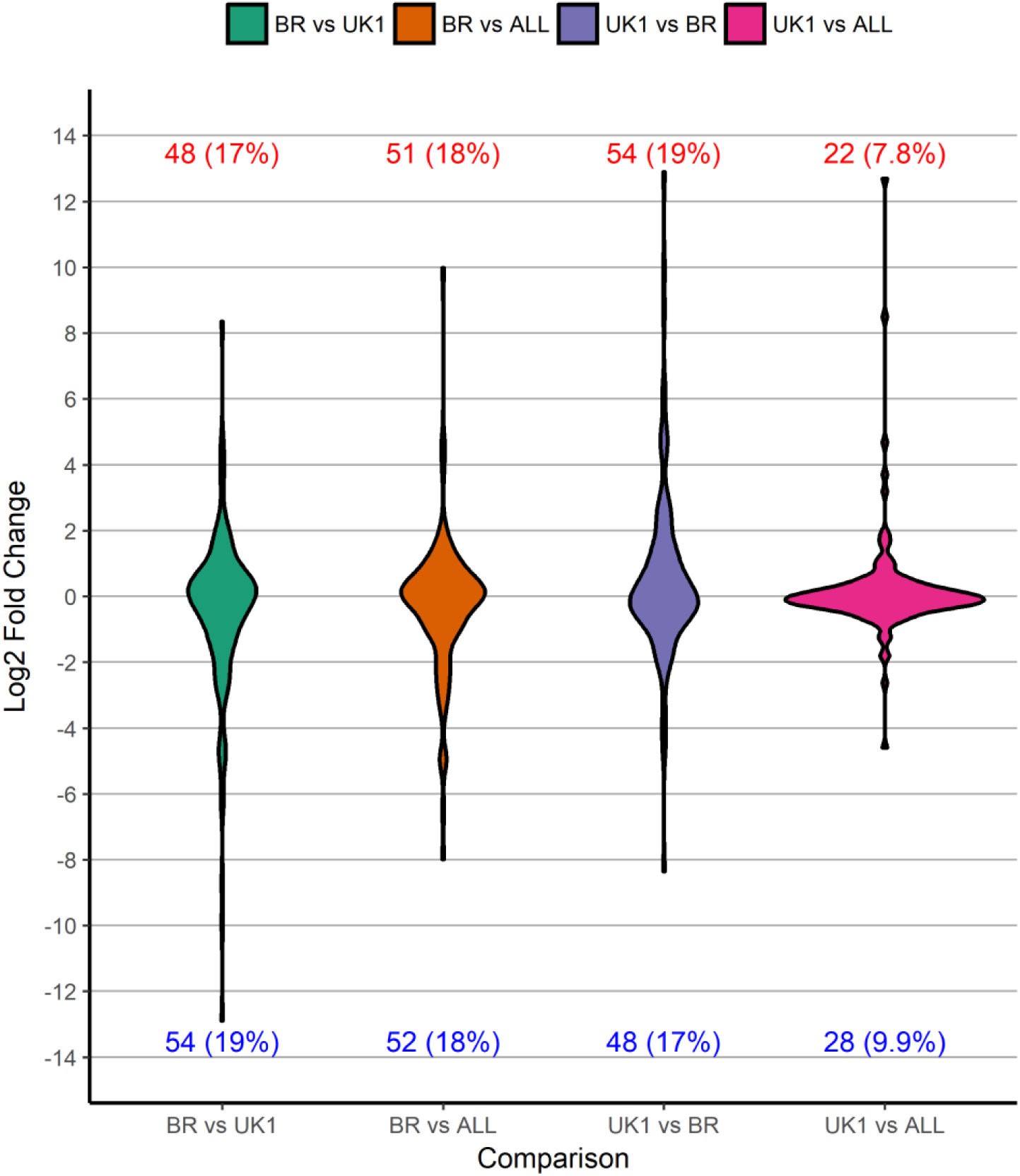
Violin plot showing significantly up-regulated and down-regulated microRNA genes across pairwise strain comparisons. Statistically significant (P<0.0001****) differentially expressed genes are plotted irrespective of log2 fold change. Total number and relative percentage of up-regulated and down-regulated microRNA genes are presented above and below each plot, respectively.

Three microRNAs (*Sc-mir-117, Sc-mir-27*, and *Sc-mir-774*) were highly differentially expressed, correlating with enhanced chemotaxis behaviour in *S. carpocapsae* Breton (>6 log2 fold, P<0.0001**** relative to UK1 and All strains). A further 18 microRNAs were likewise differentially expressed and correlated with enhanced chemotaxis behaviour (>2 log2 fold, P<0.0001****) (Fig 6A; supplemental files S3 & S4). Comparatively fewer microRNAs were differentially regulated and correlated with reduced nictation behaviour in *S. carpocapsae* UK1, with *Sc-mir-772* (up-regulated 12.9 and 12.7 log2 fold, P<0.0001**** relative to Breton and All strains, respectively) and *Sc-mir-773* (up-regulated 8.7 and 8.5 log2 fold, P<0.0001**** relative to Breton and All strains, respectively) representing notable exceptions. A further five microRNAs (*Sc-mir-754, Sc-mir-756, Sc-mir-760, Sc-let-7* and *Sc-mir-84-5pi*) were likewise differentially expressed and correlated with reduced nictation behaviour (>2 log2 fold, P<0.0001***) (Fig 6B; supplemental files S3 & S4).

**Figure 6.**
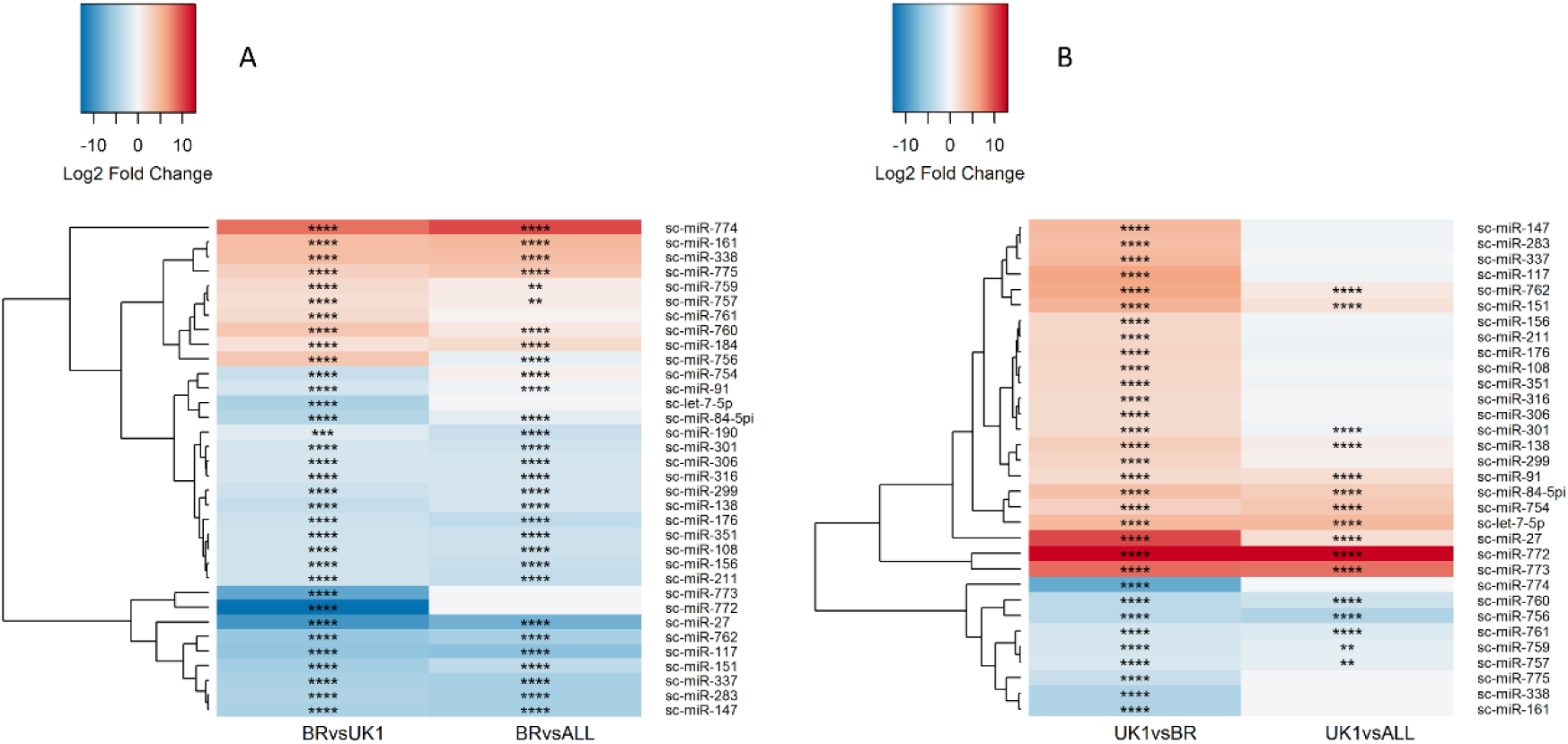
Differential expression analysis of microRNAs that correlate with *S. carpocapsae* strain behaviour. Heatmaps showing differential expression of microRNAs with at least one pairwise expression difference of >2 log2 fold change, P<0.0001****. (A) Differentially expressed microRNAs in *S. carpocapsae* Breton, relative to ALL and UK1 strains; assessing shared expression patterns that correlate with increased chemotaxis behaviour. (B) Differentially expressed microRNAs in *S. carpocapsae* UK1, relative to ALL and Breton strains; assessing shared expression patterns that correlate with reduced nictation behaviour (supplemental file S4).

### MicroRNA target analysis

*In silico* microRNA target prediction through miRanda suggests a substantial bias towards interactions with gene 5’UTRs. A total of 231,120 microRNA interactions, representing 247,237 binding sites are predicted for the 5’UTR of *S. carpocapsae* genes, relative to 127,796 microRNA interactions, representing 132,309 binding site predictions within the 3’UTR of *S. carpocapsae* genes. Through screening each of the most differentially expressed (>2 log2 fold change, P<0.0001****) microRNAs that correlate with behavioural variants across pairwise comparisons (Fig 6; supplemental file S4), we identify a substantial number of predicted interactions with neuronal gene families considered in this study (Table 1). These datasets demonstrate potential cooperation between microRNAs that are predicted to interact with shared target genes. For example, the ion channel gene *Sc-asic-2ii* is a predicted target for *Sc-mir-147, Sc-mir-301-3p*, and *Sc-mir-316*, all highly differentially expressed, and correlated with increased chemotaxis behaviour (Table 1; Fig 5A). Similarly, the *let-7* family members, *Sc-mir-84-5pi* and *Sc-let-7* co-ordinately target two *ins-1* paralogues; both microRNAs are highly differentially expressed, and correlated with reduced nictation behaviour in *S. carpocapsae* UK1 (Fig 5B). Our data also suggest that different microRNAs are predicted to interact with and converge on a number of shared neuronal gene targets considered here (Table 1). For example, *Sc-mir-301-3p* is predicted to simultaneously target the *twk-12, egas-1* and *asic-2* ion channel genes, alongside the *nlp-39* neuropeptide-like protein gene, correlating with increased chemotaxis behaviour (Table 1; Fig 5A & 1A). MiRanda microRNA target predictions across both strict and unrestricted settings for global 5’ and 3’UTRs are presented in supplemental files S5-S8.

**Table 1.**
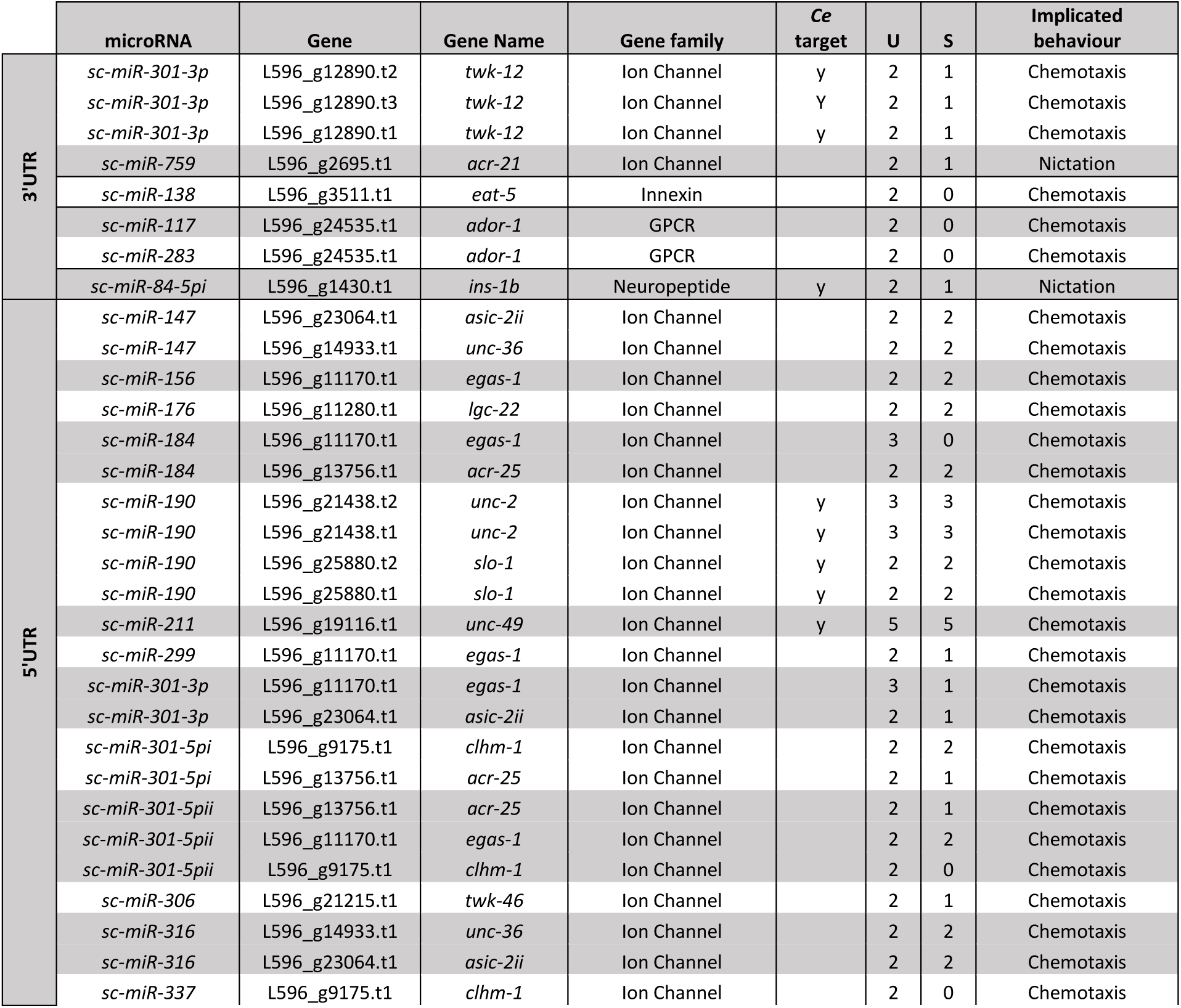
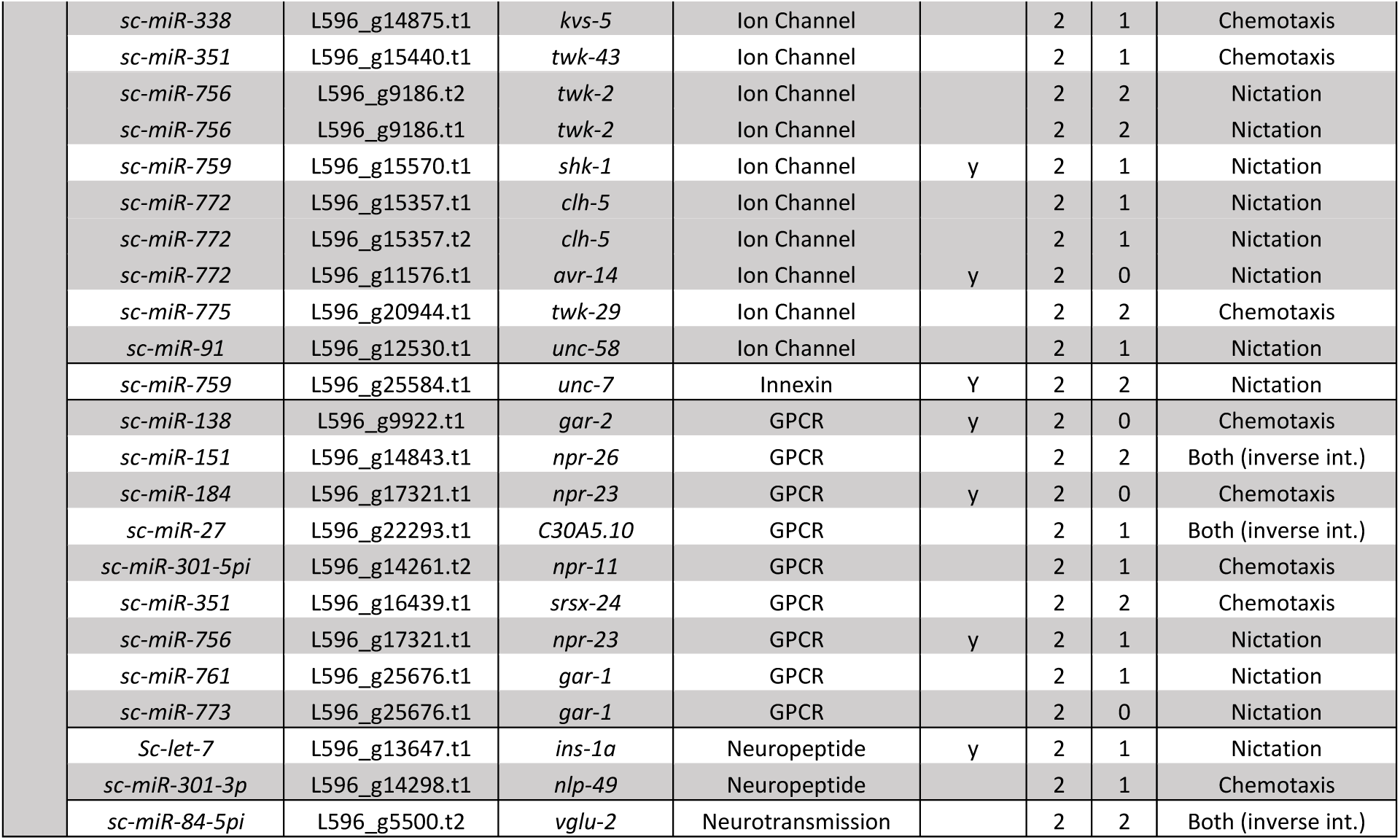
Predicted gene targets for differentially expressed microRNAs (>2 log2 fold, P<0.0001) that correlate with *S. carpocapsae* strain behaviour. Differentially expressed *S. carpocapsae* microRNAs were assessed for binding sites across 5’ and 3’UTRs of all ion channel, innexin, GPCR, neurotransmitter and neuropeptide genes. *Ce* target refers to direct 1 to 1 gene orthologues that are biochemically confirmed microRNA targets in *C. elegans* [21]. U refers to the number of predicted interacting microRNAs following unrestricted microRNA target prediction; S refers to the number of predicted interacting microRNAs following strict microRNA target prediction. Within the implicated behaviour column, “Both (inverse int.)” refers to involvement in both behaviours, through polarised regulation states (significantly upregulated for one, and significantly downregulated for the other).

|  | microRNA | Gene | Gene Name | Gene family | Ce target | U | S | Implicated behaviour |
| --- | --- | --- | --- | --- | --- | --- | --- | --- |
| 3'UTR | *sc-miR-301-3p* | L596_g12890.t2 | *twk-12* | Ion Channel | y | 2 | 1 | Chemotaxis |
|  | *sc-miR-301-3p* | L596_g12890.t3 | *twk-12* | Ion Channel | Y | 2 | 1 | Chemotaxis |
|  | *sc-miR-301-3p* | L596_g12890.t1 | *twk-12* | Ion Channel | y | 2 | 1 | Chemotaxis |
|  | *sc-miR-759* | L596_g2695.t1 | *acr-21* | Ion Channel |  | 2 | 1 | Nictation |
|  | *sc-miR-138* | L596_g3511.t1 | *eat-5* | Innexin |  | 2 | 0 | Chemotaxis |
|  | *sc-miR-117* | L596_g24535.t1 | *ador-1* | GPCR |  | 2 | 0 | Chemotaxis |
|  | *sc-miR-283* | L596_g24535.t1 | *ador-1* | GPCR |  | 2 | 0 | Chemotaxis |
|  | *sc-miR-84-5pi* | L596_g1430.t1 | *ins-1b* | Neuropeptide | y | 2 | 1 | Nictation |
| 5'UTR | *sc-miR-147* | L596_g23064.t1 | *asic-2ii* | Ion Channel |  | 2 | 2 | Chemotaxis |
|  | *sc-miR-147* | L596_g14933.t1 | *unc-36* | Ion Channel |  | 2 | 2 | Chemotaxis |
|  | *sc-miR-156* | L596_g11170.t1 | *egas-1* | Ion Channel |  | 2 | 2 | Chemotaxis |
|  | *sc-miR-176* | L596_g11280.t1 | *lgc-22* | Ion Channel |  | 2 | 2 | Chemotaxis |
|  | *sc-miR-184* | L596_g11170.t1 | *egas-1* | Ion Channel |  | 3 | 0 | Chemotaxis |
|  | *sc-miR-184* | L596_g13756.t1 | *acr-25* | Ion Channel |  | 2 | 2 | Chemotaxis |
|  | *sc-miR-190* | L596_g21438.t2 | *unc-2* | Ion Channel | y | 3 | 3 | Chemotaxis |
|  | *sc-miR-190* | L596_g21438.t1 | *unc-2* | Ion Channel | y | 3 | 3 | Chemotaxis |
|  | *sc-miR-190* | L596_g25880.t2 | *slo-1* | Ion Channel | y | 2 | 2 | Chemotaxis |
|  | *sc-miR-190* | L596_g25880.t1 | *slo-1* | Ion Channel | y | 2 | 2 | Chemotaxis |
|  | *sc-miR-211* | L596_g19116.t1 | *unc-49* | Ion Channel | y | 5 | 5 | Chemotaxis |
|  | *sc-miR-299* | L596_g11170.t1 | *egas-1* | Ion Channel |  | 2 | 1 | Chemotaxis |
|  | *sc-miR-301-3p* | L596_g11170.t1 | *egas-1* | Ion Channel |  | 3 | 1 | Chemotaxis |
|  | *sc-miR-301-3p* | L596_g23064.t1 | *asic-2ii* | Ion Channel |  | 2 | 1 | Chemotaxis |
|  | *sc-miR-301-5pi* | L596_g9175.t1 | *clhm-1* | Ion Channel |  | 2 | 2 | Chemotaxis |
|  | *sc-miR-301-5pi* | L596_g13756.t1 | *acr-25* | Ion Channel |  | 2 | 1 | Chemotaxis |
|  | *sc-miR-301-5pii* | L596_g13756.t1 | *acr-25* | Ion Channel |  | 2 | 1 | Chemotaxis |
|  | *sc-miR-301-5pii* | L596_g11170.t1 | *egas-1* | Ion Channel |  | 2 | 2 | Chemotaxis |
|  | *sc-miR-301-5pii* | L596_g9175.t1 | *clhm-1* | Ion Channel |  | 2 | 0 | Chemotaxis |
|  | *sc-miR-306* | L596_g21215.t1 | *twk-46* | Ion Channel |  | 2 | 1 | Chemotaxis |
|  | *sc-miR-316* | L596_g14933.t1 | *unc-36* | Ion Channel |  | 2 | 2 | Chemotaxis |
|  | *sc-miR-316* | L596_g23064.t1 | *asic-2ii* | Ion Channel |  | 2 | 2 | Chemotaxis |
|  | *sc-miR-337* | L596_g9175.t1 | *clhm-1* | Ion Channel |  | 2 | 0 | Chemotaxis |

|  |  |  |  |  |  |  |  |
| --- | --- | --- | --- | --- | --- | --- | --- |
| *sc-miR-338* | L596_g14875.t1 | *kvs-5* | Ion Channel |  | 2 | 1 | Chemotaxis |
| *sc-miR-351* | L596_g15440.t1 | *twk-43* | Ion Channel |  | 2 | 1 | Chemotaxis |
| *sc-miR-756* | L596_g9186.t2 | *twk-2* | Ion Channel |  | 2 | 2 | Nictation |
| *sc-miR-756* | L596_g9186.t1 | *twk-2* | Ion Channel |  | 2 | 2 | Nictation |
| *sc-miR-759* | L596_g15570.t1 | *shk-1* | Ion Channel | y | 2 | 1 | Nictation |
| *sc-miR-772* | L596_g15357.t1 | *clh-5* | Ion Channel |  | 2 | 1 | Nictation |
| *sc-miR-772* | L596_g15357.t2 | *clh-5* | Ion Channel |  | 2 | 1 | Nictation |
| *sc-miR-772* | L596_g11576.t1 | *avr-14* | Ion Channel | y | 2 | 0 | Nictation |
| *sc-miR-775* | L596_g20944.t1 | *twk-29* | Ion Channel |  | 2 | 2 | Chemotaxis |
| *sc-miR-91* | L596_g12530.t1 | *unc-58* | Ion Channel |  | 2 | 1 | Nictation |
| *sc-miR-759* | L596_g25584.t1 | *unc-7* | Innexin | Y | 2 | 2 | Nictation |
| *sc-miR-138* | L596_g9922.t1 | *gar-2* | GPCR | y | 2 | 0 | Chemotaxis |
| *sc-miR-151* | L596_g14843.t1 | *npr-26* | GPCR |  | 2 | 2 | Both (inverse int.) |
| *sc-miR-184* | L596_g17321.t1 | *npr-23* | GPCR | y | 2 | 0 | Chemotaxis |
| *sc-miR-27* | L596_g22293.t1 | *C30A5.10* | GPCR |  | 2 | 1 | Both (inverse int.) |
| *sc-miR-301-5pi* | L596_g14261.t2 | *npr-11* | GPCR |  | 2 | 1 | Chemotaxis |
| *sc-miR-351* | L596_g16439.t1 | *srsx-24* | GPCR |  | 2 | 2 | Chemotaxis |
| *sc-miR-756* | L596_g17321.t1 | *npr-23* | GPCR | y | 2 | 1 | Nictation |
| *sc-miR-761* | L596_g25676.t1 | *gar-1* | GPCR |  | 2 | 1 | Nictation |
| *sc-miR-773* | L596_g25676.t1 | *gar-1* | GPCR |  | 2 | 0 | Nictation |
| *Sc-let-7* | L596_g13647.t1 | *ins-1a* | Neuropeptide | y | 2 | 1 | Nictation |
| *sc-miR-301-3p* | L596_g14298.t1 | *nlp-49* | Neuropeptide |  | 2 | 1 | Chemotaxis |
| *sc-miR-84-5pi* | L596_g5500.t2 | *vglu-2* | Neurotransmission |  | 2 | 2 | Both (inverse int.) |

### Differential isoform usage between strains

Two genes implicated in the synthesis and transport of classical neurotransmitters were found to exhibit statistically significant differences in isoform preference across *S. carpocapsae* strains following RSEM analysis (Fig 7 & 8; Supplemental file S9). Interestingly, these isoform variants exhibit altered UTR sequences in additional to altered exon usage. *In silico* microRNA target predictions were conducted for all predicted microRNAs relative to the respective isoform UTRs, identifying quantitative differences in predicted microRNA interactions with the various gene isoforms (Table 2; Supplemental file S9).

**Figure 7.**
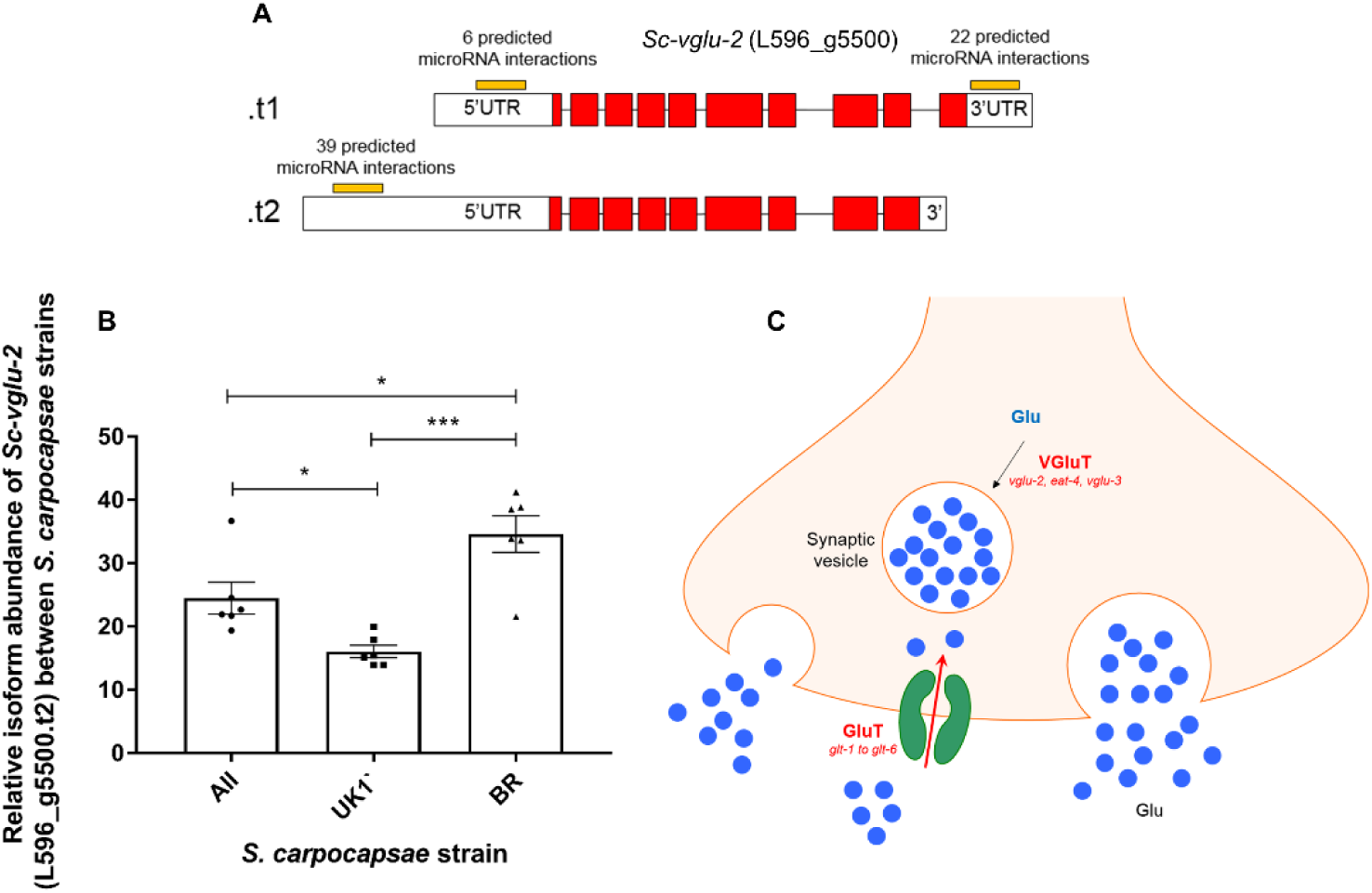
Variation in isoform preference may alter microRNA targeting of the vesicular glutamine transporter gene *Sc-vglu-2* between strains of *S. carpocapsae*. **(A)** Diagrammatic depiction of isoform structure between predicted variants (not to scale); white boxes indicate UTRs, red boxes indicate exons. **(B)** Graph indicating relative percentage abundance of L596_g5500.t2 across *S. carpocapsae* strains (n=6 independent libraries per strain). One way ANOVA and Tukey’s multiple comparison tests were conducted using Graphpad Prism 7.02. P<0.05*, P<0.001***. **(C)** Diagrammatic depiction of the relative position of VGluT in a glutamatergic neuron.

**Figure 8.**
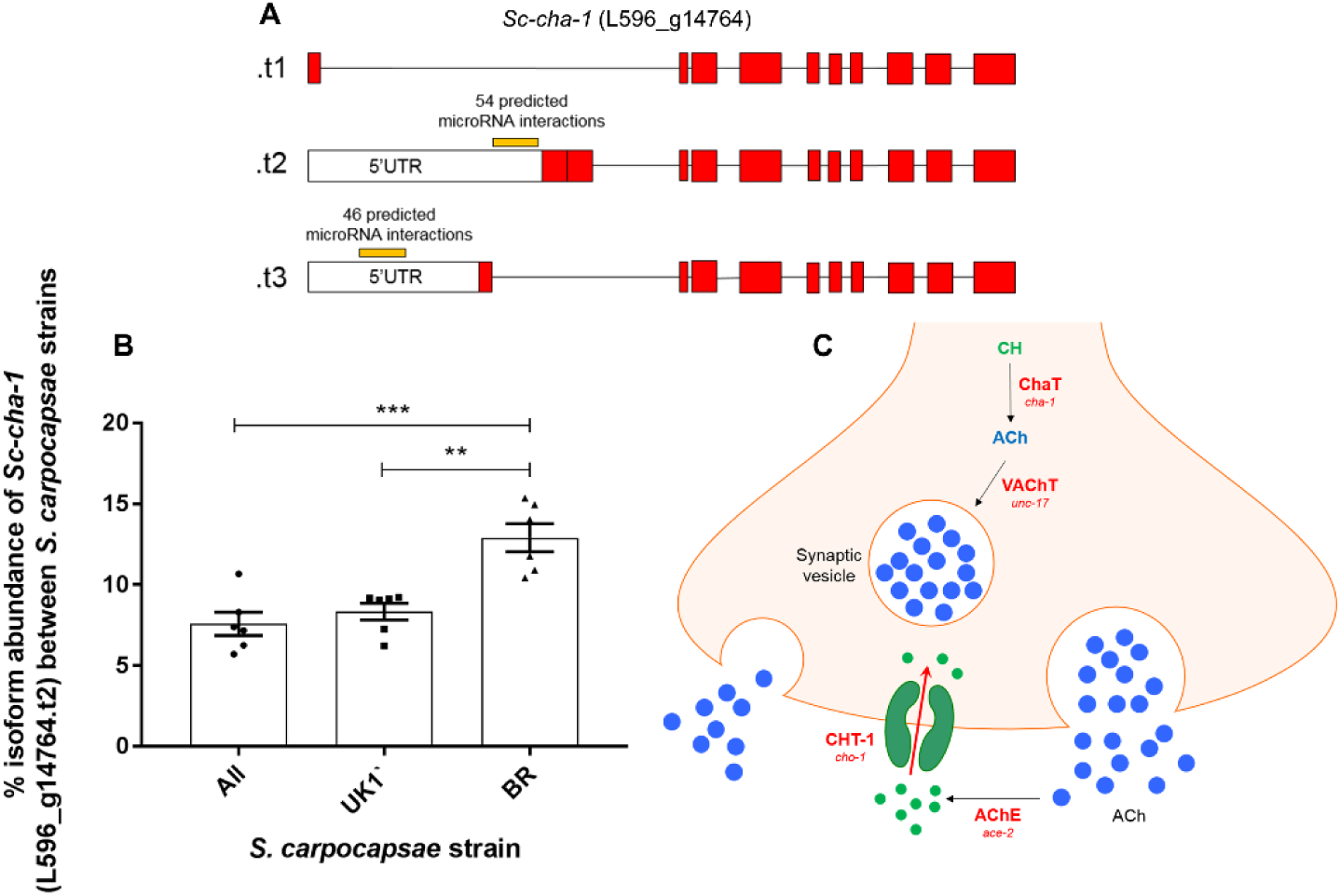
Variation in isoform preference may alter microRNA targeting of the choline acetyltransferase gene *Sc-cha-1* between strains of *S. carpocapsae*. **(A)** Diagrammatic depiction of isoform structure between predicted variants (not to scale); white boxes indicate UTRs, red boxes indicate exons. **(B)** Graph indicating relative percentage abundance of L596_g14764.t2 across *S. carpocapsae* strains (n=6 independent libraries per strain). One way ANOVA and Tukey’s multiple comparison tests were conducted using Graphpad Prism 7.02. P<0.01**, P<0.001***. (C) Diagrammatic depiction of the relative position of CHaT in a cholinergic neuron.

**Table 2.** Differential microRNA targeting across *Sc-cha-1* and *Sc-vglu-2* gene isoform 5’ and 3’UTRs. U denotes number of predicted microRNA interactions following unrestricted target prediction in miRanda; S denotes number of predicted microRNA interactions following strict target prediction in miRanda. Numbers in brackets indicate the relative number of binding sites across both U and S settings.

| Transcript ID | Mean %Isoform abundance |  |  | N° microRNA interactions in 3'UTR (binding sites) |  | N° microRNA interactions in 5'UTR (binding sites) |  |
| --- | --- | --- | --- | --- | --- | --- | --- |
|  | ALL | UK1 | BR | U | S | U | S |
| L596_g14764.t1 | 90.9% | 90.1% | 85.8% | - | - | - | - |
| L596_g14764.t2 | 7.6% | 8.3% | 12.9% | - | - | 54 (57) | 36 (36) |
| L596_g14764.t3 | 1.5% | 1.6% | 1.3% | - | - | 46 (49) | 31 (31) |
| L596_g5500.t1 | 75.5% | 83.9% | 65.4% | 22 (22) | 5 (5) | 6 | 4 (4) |
| L596_g5500.t2 | 24.5% | 16.1% | 34.6% | - | - | 39 (49) | 25 (27) |

## Discussion

Understanding the link between genotype and phenotype has been a long-standing preoccupation of the life sciences. Various approaches can be used to link gene sequence to function, from high resolution study of individual genes through forward and reverse genetics, to genome and transcriptome-wide associative studies that correlate sequence or abundance differences with phenotypic variants. In this study our aim was to use a small scale, high resolution, associative transcriptomics approach to correlate protein-coding and non-coding RNA expression patterns with distinct IJ behavioural states across *S. carpocapsae* strains. Relative to genome-wide associative studies (GWAS), associative transcriptomics can provide a more obvious functional linkage between genotypic differences, and functional consequences, through the identification of differentially expressed genes and isoform variants.

Our approach has generated high quality transcriptomes (n=18) and small RNA libraries (n=18) for the IJ stage of three behaviourally distinct *S. carpocapsae* strains, and has catalogued a number of protein-coding and non-coding RNAs that correlate strongly with distinct host-finding behaviours. We have applied an arbitrary threshold of >2 log2 fold change (P<0.0001****) across both pairwise strain comparisons in order to link gene expression markers to behavioural differences. This is a stringent purifying criteria that selects for the most highly differentially expressed genes, which are most likely to have biologically relevant impacts on function and behaviour. Many of these genes have already been shown to regulate behaviour in the model nematode *C. elegans*, however none have been implicated specifically in host-finding behaviours of parasitic nematodes, pointing to the discovery potential of this approach. Ultimately, an associative transcriptomic study will be strengthened by assaying a wide range of specific and reproducible phenotypes, across a wide range of comparator strains. This proof of concept study demonstrates the potential benefit of establishing and investing in natural diversity resources for parasitic nematodes, alongside more expansive studies in the model nematode *C. elegans* [50, 51]. To that end, we are committed to establishing such a resource for *Steinernema* spp. that will grow to include genomic, transcriptomic, small RNA and behavioural metadata across a wide range of curated and maintained strains. The *S. carpocapsae* strains used in this study could be used for additional behavioural and phenotypic studies, the outputs of which could then be correlated with the transcriptomes and small RNA libraries generated here, to give gene expression markers and/or isoform variant patterns that associate with the phenotype or behaviour of interest.

Our data point to the potential involvement of several different neuronal gene families in the regulation of host-finding behaviours. Only one neuropeptide gene, *nlp-36*, exhibits co-ordinate differential expression (>2 log2fold, P<0.0001****) that correlates with distinct strain behaviours. NLP-36 is positively regulated by the cyclic nucleotide-gated channel subunit TAX-2 in *C. elegans*, which is implicated in the regulation of olfaction, chemosensation, thermosensation and axon guidance for a number of sensory neurons [52, 53]. In *S. carpocapsae* UK1, significant co-ordinate up-regulation of *nlp-36* correlates with reduced nictation behaviour (Fig 1B & 3B). Down-regulation of insulin signalling during *C. elegans* development has been found to influence nictation behaviour in dauer juveniles, indicating a tuning of behaviour to environmental conditions [11]. Gruner et al. [54] point to a potential neuronal programming circuit for behavioural adjustment in *C. elegans* through the combined influence of insulin signalling, TRPV signalling and agonism of NPR-1, a known receptor of FLP-21 and FLP-18. Although the differential expression of *daf-28* does not meet the shared >2 log2 fold change threshold we have applied to sequencing datasets, it is nonetheless down-regulated in *S. carpocapsae* Breton, relative to the other strains, correlating with enhanced chemotaxis toward *G. mellonella.* Conversely, *daf-28* was up-regulated in UK1 relative to both Breton and All strains (Fig 3A-B), correlating with reduced nictation (Fig 1B). As was found for the two *asic-2* sodium channel paralogues, *daf-28* and other genes could function as part of a neurobiological switch, tuning behavioural strategy toward either migratory or stationary host-finding modes. It may be informative to track the dynamic abundance of such genes in IJs naturally enacting and transitioning between these behaviours, or in conditions that are known to enhance or suppress these behaviours to further strengthen the correlation of expression with behaviour [55, 56].

The up-regulation of srsx GPCR genes in *S. carpocapsae* Breton correlates with enhanced chemotaxis to the lab host insect *G. mellonella* (Fig 4A). One of these genes is orthologous to a cluster of srsx genes (*srsx-22* and *srsx-24*) that are enriched in dauer stage *C. elegans.* It has been shown that dauer stage *Caenorhabditis* spp. are attracted to certain insect species, increasing the opportunity to engage in phoresis [57]. This cluster of shared srsx GPCRs may therefore mediate this attraction, in isolation, or in synergy with other such receptors. Two *asic-2* sodium channel paralogues (denoted here as *Sc-asic-2i* and *Sc-asic-2ii*) are also notable as being differentially expressed and inversely correlated with both increased chemotaxis and reduced nictation behaviours (Fig 4E-F). ASIC-2 is known to regulate aspects of nematode body posture and mechanosensation in *C. elegans* [58].

Small non-coding RNAs, including microRNAs, are increasingly implicated in complex aspects of biology and behaviour [51, 59–61]. We conducted a deep analysis of small RNA profiles across *S. carpocapsae* strains, and found a substantial degree of variation in relative abundance (Figs 5 & 6, supplemental file S3). Numerous microRNAs are differentially expressed, and correlate with behavioural differences across pairwise comparisons (Fig 6, supplemental file S4). Key to extrapolating biologically relevant information from microRNA networks is the identification of gene targets. Although there are many *in silico* tools that predict microRNA-mRNA transcript interactions, false positives are likely to be common [62]. Whilst many factors influence the reality and significance of predicted interactions, bioavailability of target gene and microRNA in terms of special and temporal expression patterns will be key, along with the number of available microRNA binding sites, the relative enthalpy of binding interactions, and local competition for available microRNAs. In order to build on the basic knowledge presented in this study, it will be necessary to biochemically validate gene transcripts as microRNA targets through Argonaute ClIP-seq [46], and further, to demonstrate co-localisation of microRNA and target mRNA transcript to confirm interaction of discrete partners. This represents a substantial, but necessary task if we are to unravel the biological significance of microRNA regulation in the context of complex phenotypes and behaviours. Previous publications have employed a hierarchical and cooperative *in silico* target prediction approach using several programs simultaneously to arrive at an agreed set of targets [59]. Whilst this will certainly reduce the complexity of any target gene set, it will also constrain the output according to the most stringent program, leading inevitably to false negatives when perfect seed site complementarity is required by one or more programs. Here we have employed a dual analysis strategy within the miRanda discovery environment [44], using both unrestricted and strict discovery modes; the latter requires perfect seed site complementarity between microRNA and target mRNA transcript. Collectively, this strategy maps well to biochemically-validated microRNA target interactions, including those that do not require perfect seed site binding [46]. However, as with any *in silico* prediction approach, biological validation is still required to corroborate interactions.

Unexpectedly, our datasets suggest a strong microRNA target site enrichment within predicted 5’UTRs of *S. carpocapsae*. This may be biological significant, or could represent an artefact of computational UTR prediction. Ultimately, higher confidence interactions could be established by sequencing full length transcripts and confirming UTR identity on a transcriptome wide scale. *In silico* target prediction for differentially expressed (>2 log2 fold,P<0.0001****) and behaviourally correlated microRNAs points to instances of potential microRNA co-operation. For example, the ion channel genes *Sc-asic-2ii* and *Sc-egas-1* are predicted targets for three and five individual differentially expressed microRNAs each (Table 1). The neuropeptide GPCR gene, *Sc-npr-23* is likewise targeted independently by two differentially expressed microRNAs. The differential expression pattern of each targeting microRNA correlates with altered chemotaxis behaviour, suggesting that microRNAs may also drive behavioural variation (Table 1).

The predicted targeting of *Sc-npr-11* by *sc-miR-301-5pi* reveals that UTR sequence variation between the two annotated *npr-11* isoforms allows the dominant isoform (representing ∼75% of all *npr-11* transcript across strains) to escape *mir-301-5pi* interaction (Table 1; supplemental file S9). Likewise, variation in the 5’UTR and 3’UTRs of *Sc-vglu-2* and *Sc-cha-1* (Figs 6 & 7, Table 2) reveal a potential quantitative difference in microRNA interaction events across different gene isoforms. In the case of *Sc-cha-1*, gene-level regulation of microRNA target site visibility could represent a strain specific mechanism to dampen global cholinergic signalling, or could allow selective dampening in a subset of cholinergic neurons. Ascertaining the relevance of isoform variance as it relates to microRNA interactions will require detailed gene localisation and functional studies, but represents a fascinating area that is presently under-studied. Our datasets highlight 5’ and 3’UTR variation as a factor in differential microRNA target site visibility, although it seems especially likely that alternative polyadenylation signals in the 3’UTR of protein-coding genes will exert substantial control over microRNA target site visibility, especially given the evident pervasiveness of 3’UTR variation in nematodes [63, 64]. Any such regulation could modify the relative percentage of gene isoform variants accessible to microRNAs, which could allow for cell, or tissue-specific regulatory events that may be difficult to ascertain from organism-wide datasets. Collectively, our data point to a complex co-regulatory environment involving gene-specific isoform variation, and microRNA transcriptional regulation that is likely to influence various aspects of biology, including behaviour.

In the same way that “the best model for a cat is several cats” [65], the best model for a nematode parasite of vertebrates, is a sustainable population of the very same. However, this might not always be preferable in terms of ethics, available tools, sustainability or reproducibility. ‘Model’ status for any organism is contingent on high quality genomic and transcriptomic resources, easily amenable research tools, both in terms of genetic and molecular manipulation, alongside robust behavioural and phenotypic assays. Ease of culture, handling, and short generation times should also be major considerations. A model organism and the data generated from it must also be sufficiently relevant to trigger near-term impact on other species that have implications for health and economy. The datasets presented in this study build upon a growing catalogue of tools and resources for *Steinernema* spp. entomopathogenic nematodes [10, 22, 31, 66–68]. The close phylogenetic relationship between *Steinernema* spp. and economically important mammalian parasites [69], coupled with striking behavioural similarities, and a pathogenic life style that can be fully recapitulated in a Petri dish suggests that these entomopathogenic nematodes could represent attractive and convenient new surrogate models for parasite biology and behaviour. In addition to their potential worth as model organisms, *Steinernema* spp. have attracted considerable attention as bioinsecticidal agents [70]. The interest in *Steinernema* spp. as biological models, and as economically relevant endpoint organisms in their own right represents a unique proposition for researchers interested in comparative biology, behaviour and host-parasite interactions.

## Acknowledgements

Waxworms infected with *S. carpocapsae* (Breton), were a gift from Prof Nelson Simões, University of the Azores, Portugal. *S. carpocapsae* (UK1) IJs were a gift from BASF. Microdirt chip PDMS moulds were a gift from Prof Junho Lee, Seoul National University, Korea. The authors would like to thank Matthew Sturrock and Steven Dyer for critical reading of the manuscript.

## Biological resources and dataset availability

The *S. carpocapsae* strains used in this study are maintained within the Dalzell lab, and are freely available upon request, as are other local isolates of *Steinernema feltiae*, collected from the UK and Ireland. We would be grateful to receive additional *Steinernema* spp. strains from the research community in order to develop a natural diversity resource, which will be free to access and build upon. Illumina HiSeq reads for transcriptome and small RNAs across each *S. carpocapsae* strain are available within the SRA library under project code: PRJNA436466.

## Supporting information captions

**Supplemental file S1**. List of all differentially expressed genes (>2 log2 fold, P<0.0001) across all pairwise strain comparisons.

**Supplemental file S2.** List of all differentially expressed genes represented in heatmaps, including annotated gene names, fold changes across pairwise comparisons, and adjusted P values.

**Supplemental file S3.** List of *S. carpocapsae* microRNAs, sequences and read counts across strains. **Supplemental file S4.** List of differentially expressed microRNAs (>2 log2 fold, P<0.0001), including pairwise fold changes, and adjusted P values.

**Supplemental file S5.** MicroRNA target predictions 1: Miranda output file for microRNA targets in 5’UTRs using unrestricted setting.

**Supplemental file S6.** MicroRNA target predictions 2: Miranda output file for microRNA targets in 5’UTRs using strict setting.

**Supplemental file S7.** MicroRNA target predictions 3: Miranda output file for microRNA targets in 3’UTRs using unrestricted setting.

**Supplemental file S8.** MicroRNA target predictions 4: Miranda output file for microRNA targets in 3’UTRs using stricted setting.

**Supplemental file S9.** RSEM isoform quantification file for *S. carpocapsae* strains.

